# Genome-wide analysis of DNA methylation reveals selection signatures of grass carp during the domestication

**DOI:** 10.1101/2021.11.04.467282

**Authors:** Yichao Li, Bing Fu, Junming Zhang, Jun Xie, Guangjun Wang, Peng Jiang, Jingjing Tian, Hongyan Li, Yun Xia, Ermeng Yu

**Affiliations:** Pearl River Fisheries Research Institute, Chinese Academy of Fishery Sciences, Guangzhou 510380, China

**Keywords:** DNA methylation, Selection signatures, Grass carp, Domestication, Epigenetics

## Abstract

With the rapid development of aquaculture, more and more fish species from wild environments are artificially domesticated and cultured. In the process of domestication, the fish develop some adaptations and phenotypic traits, namely selection signatures. However, it is still unclear about the biological process underlying these selection signatures. Here, we used grass carp (*Ctenopharyngodon idellus*), an aquaculture fish with the largest production worldwide, to detect its selection signatures and investigate the roles of DNA methylation in the emergence of selection signatures during domestication based on whole-genome bisulfite sequencing technology. Our results showed that domesticated grass carp demonstrated four selection signatures, including growth and metabolism, immunity, foraging and learning behaviors, and 38 candidate genes were associated with these traits. 16 of candidate genes, such as IGF-1, GK, GYS1, etc., were found to play major roles in the growth and metabolism. Immunity signature was related to 11 of candidate genes, including MHCI, MHCII, C1QA, etc. The GRM1, TAS1R1 and TAS1R3 genes were essential for the adaptation of domesticated grass carp to commercial feed in artificial rearing condition. The C-FOS, POMC and CBP genes might be responsible for the acquisition of novel feeding habits and contribute to faster growth indirectly by enhancing food intake. These findings would provide new insights to expand our understanding on the role of DNA methylation in shaping physiological phenotypes in fish, and also contribute to efficient breeding of aquaculture stocks and restocking programs.

## 1. Introduction

As an economically food source of humans, fish is one of the most important domesticated species. In modern agricultural industry, fish have undergone strong long-term artificial selection and developed a range of adaptations and phenotypic traits, namely selection signatures (Milla et al., 2021). Fish selection signatures usually include the higher flexibility in diet, rapid growth, less stress susceptibility, a more socially tolerant disposition and enhanced prolificacy (Pasquet et al., 2018), and they distinguish the domesticated breeds from their wild counterparts. These phenotypic differences make fish highly suitable for animal agriculture and comparative studies.

Epigenetic changes is increasingly recognized to contribute to the emergence of phenotypic differences (Goldberg et al., 2007). DNA methylation, the most widely studied epigenetic regulatory mechanisms, can respond to environmental changes and, at the same time, be stable enough to be maintained throughout lifetime, even across generations (Anastasiadi et al., 2019). DNA methylation has been associated with many biological process, including gene expression regulation, genomic imprinting, X chromosome inactivation, embryonic development, the alteration of chromatin structure and transposon inactivation, etc. (Jones et al., 2012). Compared to other approaches which have been developed to analyze DNA methylation profiles at the genome-wide levels, whole-genome bisulfite sequencing (WGBS) has two major advantages of assessing the methylation state of nearly every CpG site and determining absolute DNA methylation levels (Yong et al., 2016). WGBS has been broadly employed to analyze the genome-wide methylation profiles of many animals, including sea cucumber (Yang et al., 2020) and Chinese perch (Pan et al., 2021).

In recent years, some studies explore the roles of DNA methylation regulation on fish during domestication. However, these studies mainly focus on the role of DNA methylation in the changes of one particular phenotypic characteristics, such as growth (Li et al., 2017), reproduction (Podgorniak et al., 2019), morphological changes (Zhang et al., 2017), and learning behaviors (Dou et al., 2018). But, until now, no systematic studies have investigated the roles of DNA methylation in the emergence of different selection signatures in the process of fish domestication, such as growth and metabolism, immunity, foraging and learning behaviors.

Grass carp (*Ctenopharyngodon idellus*) is one of the most important freshwater aquaculture species, and its global production reaches 5.704 million tons in 2018, the highest in fish production worldwide, providing low-cost, high-quality animal protein especially for developing and underdeveloped regions (FAO, 2020). Although some farmers have carried out breeding for several generations, many farmers still rely on wild-caught fish for broodstock to maintain genetic diversity (Fu et al., 2015). Furthermore, farmed grass carp shares some of the characteristics of the domestication syndrome, including the changes in morphological features (Zhao et al., 2020), rapid growth (Ashraf et al., 2011) and anti-predator behavior (Tang et al, 2017), indicating that this species could be a suitable model species for studying the roles of DNA methylation in the formation of selection signatures in fish species.

This study used WGBS (whole-genome bisulfite sequencing) to carry out genome-wide DNA methylation analysis of the domesticated and wild grass carp. We obtained comprehensive DNA methylation profiles for the two groups and identified differentially methylated genes (DMGs) that might contribute to the emergence of four selection signatures differences: growth and metabolism, immunity, foraging and learning behaviors. These findings would provide new insights to expand our understanding about the role of epigenetic modifications in shaping physiological phenotypes in fish.

## 2. Materials and Methods

### 2.1 Domesticated trial

A total of 20 wild sub-adult grass carp at the approximately 18-month old determined by annulus characteristics of scale (Cai et al., 2020), were captured using traps and gill nets along the Xijiang river in Zhaoqing City, Guangdong Province, China, within a 1000 m radius from the following coordinates (latitude: 23.1034.8 °N; longitude: 112.4554.0 °E). Subsequently, the fish were transported to breeding base and randomly allocated into two ponds (1.5 m × 1.5 m × 1 m), 10 individuals in each pond for subsequent domesticated experiments in the Pearl River Fisheries Research Institute. The fish were fed commercial feed at 8:00 am and 4:00 pm each day for 180 days. The water temperature was kept at 22~27 °C, pH was 6.5~7.5, and dissolved oxygen was above 5.0 mg/L.

At the first week, the food intake of wild sub-adult grass carp was extremely low. Subsequently, some wild grass carp started to snatch food during feeding with the feed amount for each day was 0.5~2% of fish weight. After about two month training, the behavior of competition for food was obvious, and the feed amount for each day remained 2~3% of fish weight.

### 2.2 Sample Collection and Preparation

Three largest domesticated grass carp after six months of domestication were collected from ponds (with a body weight of 680 ± 25 g). Wild grass carp at the approximately 24-month old were collected from same place where wild sub-adult grass carp were captured and weighed to get a body weight of 550 ± 41 g. Then, whole blood was collected from were collected from the domesticated grass carp (sample DGC1-DGC3) and wild grass carp (sample WGC1-WGC3) groups. Subsequently, blood samples were treated with EDTAK2 and then centrifuged at 12,000 rpms for 3 min to separate red blood cells (RBCs) from serum.

The experimental protocols used in this study were approved by the Laboratory Animal Ethics Committee of Pearl River Fisheries Research Institute, CAFS, China, under permit number LAEC-PRFRI-2021-06-03.

### 2.3 Serum immune parameter analysis

Blood samples were centrifuged at 3000 g for 15 min at 4 °C and serum was stored at −80 °C for analysis. The activities of lysozyme was determined by the Ultra-Sensitive Fish ELISA Kits (Sino, China) (Kit No. YX-E21980F). Besides, the Ultra-Sensitive Fish ELISA Kits were used to measure the contents of complement C3 (Kit No. YX-E21980F) and total protein (Kit No. YX-E21980F).

### 2.4 Genomic DNA Extraction and Whole-Genome Bisulfite Sequencing

Blood is the most common source of biomarkers and materials for genetic studies, because it interacts with all organs and is easily collected. Global analysis of methylation profiles using blood DNA has been broadly used to explain phenotypic differences in growth, metabolism, immunity, learning and memory in human and other animals (Wang et al., 2017; Desrivieres et al., 2021). Thus, in this present study, genomic DNA was extracted from RBCs of domesticated grass carp and wild grass carp groups and sent to BGI (BGI Tech Co., Ltd., Shenzhen, China) for whole-genome bisulfite sequencing. For normal WGBS library constructing, the DNA was fragmented by sonication using a Bioruptor (Diagenode, Belgium) to a mean size of approximately 250 bp, followed by the blunt-ending, dA addition to 3’-end, finally, adaptor ligation(in this case of methylated adaptors to protect from bisulfite conversion), essentially according to the manufacturer’s instructions. Ligated DNA was bisulfite converted using the EZ DNA Methylation-Gold kit (ZYMO). Different Insertsize fragments were excised from the same lane of a 2% TAE agarose gel. Products were purified by using QIAquick Gel Extraction kit (Qiagen) and amplified by PCR. At last, Sequencing was performed using the HighSeq4000 or other Illumina platforms.

### 2.5 Data Filtering and Reads Alignment

After sequencing data was delivered, the raw reads were filtered by removing adaptor sequences, contamination and low-quality reads from raw reads. Low-quality reads include two types, and the reads meet any one of the two conditions will be removed: (1) Unknown bases are more than10%; (2) The ratio of bases whose quality was less than 20was over 10%. After filtering, the Clean data was then mapped to the reference genome of grass carp (Wang et al., 2015) by BSMAP, and then remove the duplication reads and merge the mapping results according to each library. The BSMAP script was BSMAP -a filename_1.clean.fq.gz -b filename_2.clean.fq.gz -o filename.sam - d ref.fa -u -v 8 -z 33 -p4 -n 0 -w20 -s 16 -f 10 -L 100. The sam files were converted to bam files using scripts (samtools view -S-b-o filename.bam filename.sam; samtools sort-m 2000000000 filename.bam filename. sort; samtools index filename. sort.bam). The mapping rate and bisulfite conversion rate of each sample were calculated.

### 2.6 Identification of Differentially Methylated Regions

The methylation level was determined by dividing the number of reads covering each mC by the total reads covering that cytosine (Lister, 2009), which was also equal the mC/C ratio at each reference cytosine (Xiang et al., 2010). The formula is Rmaverge = Nmall/Nmall+Nnmall. Nm represents the reads number of mC, while Nnm represents the reads number of non-methylation reads. Putative DMRs were identified by comparison of the sample DGC and sample WGC methylomes using windows that contained at least 5 CpG (CHG or CHH) sites with a 2-fold change in methylation level and Fisher test *p* value < = 0.05.

### 2.7 Gene Ontology and Pathway Enrichment of DMRs

GO enrichment analysis provides all GO terms that significantly enriched in a list of differentially methylated genes, comparing to a genome background, and filter the differentially methylated genes that correspond to specific biological functions. This method firstly maps all differentially methylated genes to GO terms in the database (http://www.geneontology.org/), calculating gene numbers for every term, then uses hypergeometric test to find significantly enriched GO terms in the input list of differentially methylated genes, based on ‘ GO:: TermFinder’ (http://www.yeastgenome.org/help/analyze/go-term-finder). KEGG pathway enrichment analysis helps to identify significantly enriched metabolic pathways or signal transduction pathways in differentially methylated genes comparing with the whole genome background. The calculating formula is the same as that in GO analysis.

### 2.8 Protein-protein interaction network construction based on differentially expressed mRNAs

Differentially methylated genes were selected for the construction of protein-protein interaction network. Briefly, potential or confirmed protein interactions were generated and analyzed using the STRING Version 11.0 (https://string-db.org/) based on the zebrafish database, with combined score > 0.4. The PPI network was then visualized using Cytoscape v. 3.8.2 software.

### 2.9. Data analysis

SPSS software (version 20.0) was applied for statistical analysis. All values were expressed as mean ± SE. Data were analyzed by using the Student *t*-test. The *P* value less than 0.05 was considered to be statistically significant.

## 3. Results

### 3.1 Serum immune parameters

The levels of serum lysozyme, total protein in domesticated grass carp group were significantly lower than those of wild grass carp group, indicated decreased immunity in domesticated grass carp group (Table 1). The content of serum complement C3 also exhibited a downward tendency in the domesticated grass carp group, but no significant difference was observed between two groups (Table 1).

**Table 1.**
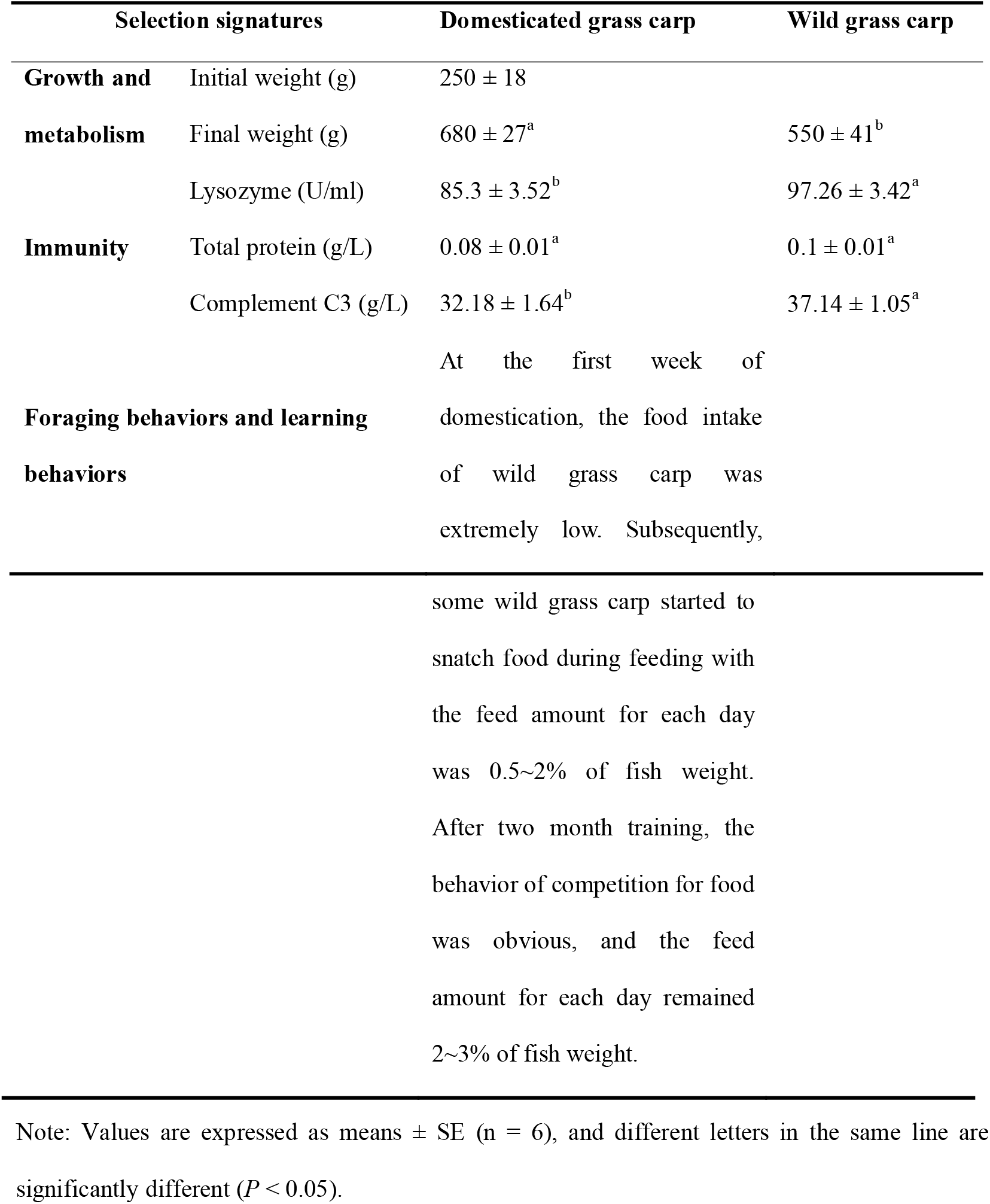
The four selection signatures of domesticated and wild grass cap

### 3.2 Global mapping and statistical analysis of the WGBS reads

We conducted whole-genome bisulfite sequencing of grass carp blood from domesticated grass carp group and wild grass carp group. After quality control of filtering, a total of 2.36 billion clean reads were generated, consisting of 378.12 million, 387.63 million, and 321.7 million reads for each domesticated grass carp sample and 371.48 million, 493.82 million, and 410.42 million reads for each wild grass carp sample (Table 2). The bisulfite conversion rate (%) of all sequencing libraries ranges from 99.16% to 99.28%. After read alignment, clean reads were mapped to the reference genome of grass carp with mapping rates ranging from 88.03 % to 92.11 % (Table 2).

**Table 2.** Alignment statistics with reference genome

| Sample ID | Clean reads | Mapped reads | Mapping rate (%) | Uniquely mapped reads | Uniquely mapping rate (%) | Bisulfite conversion rate (%) |
| --- | --- | --- | --- | --- | --- | --- |
| DGC1 | 378,121,760 | 347,944,087 | 92.02 | 329,583,800 | 87.16 | 99.28 |
| DGC2 | 387,627,636 | 353,434,830 | 91.18 | 335,393,994 | 86.52 | 99.23 |
| DGC3 | 321,704,534 | 296,306,210 | 92.11 | 281,433,956 | 87.48 | 99.16 |
| WGC1 | 371,478,196 | 321,457,393 | 86.53 | 300,678,599 | 80.94 | 99.16 |
| WGC2 | 493,819,092 | 425,920,306 | 86.25 | 398,434,532 | 80.68 | 99.24 |
| WGC3 | 410,419,590 | 361,301,972 | 88.03 | 339,621,985 | 82.75 | 99.22 |

Note: DGC, domesticated grass carp; WGC, wild grass carp.

### 3.3 Global DNA methylation patterns of domesticated and wild grass carp

The methylation levels of the whole genom were listed in Table 3. In each group, approximately 9% of all genomic C sites were methylated. Methylation in grass carp was found to exist in three sequence contexts: CG, CHG (where H is A, C, or T), and CHH. The average methylation levels of CG, CHG, and CHH at the whole genome levels were 80.3%, 1.09%, and 1.08% in the domesticated grass carp group, and 75.04%, 1.07%, and 1.13% in the wild grass carp group. The average methylation of CG showed a significantly increased within genome in the domesticated grass carp group compared with the wild grass carp group (*p* < 0.05). The methylation level of CHG and CHH showed no differences between domesticated grass carp and wild grass carp groups (*p* >0.05).

**Table 3.** Average methylation levels of different genomic regions

| Groups | Regions | C (%) | CG (%) | CHG (%) | CHH (%) |
| --- | --- | --- | --- | --- | --- |
| DGC1 | Genome | 9.76 | 80.38 | 0.99 | 1.01 |
| DGC2 | Genome | 9.69 | 80.55 | 1.18 | 1.06 |
| DGC3 | Genome | 9.68 | 79.98 | 1.11 | 1.17 |
| WGC1 | Genome | 9.21 | 74.7 | 1.12 | 1.18 |
| WGC2 | Genome | 9.19 | 75.04 | 1.02 | 1.09 |
| WGC3 | Genome | 9.13 | 75.39 | 1.07 | 1.12 |

Note: DGC, domesticated grass carp; WGC, wild grass carp. CG, CHG, and CHH (Where H is A,
C, or T).

Methylation status of CG, CHG and CHH of grass carp genome showed the methylome’s overall characteristics (Fig.1). The methylation levels of approximately 25% of all mCG were hypermethylated (methylation level > 90%). However, only about 3% of mCHH and mCHG were hypermethylated (methylation level > 90%) compared with mCG. The methylated Cs mostly occur in the form of mCG, followed by mCHH and mCHG. The methylation level distribution of mC and mCG were alike. The methylated Cs mostly occur in the form of mCG; approximately 96% of all detected mCs (Table 4). Proportion of mCHG range from 0.75% to 1.15%, and proportion of mCHH range from 2.7% to 3.05% (Table 4). The proportion of mCG, mCHG and mCHH showed no significant difference between domesticated grass carp group and wild grass carp group (*p* > 0.05).

**Fig.1.**
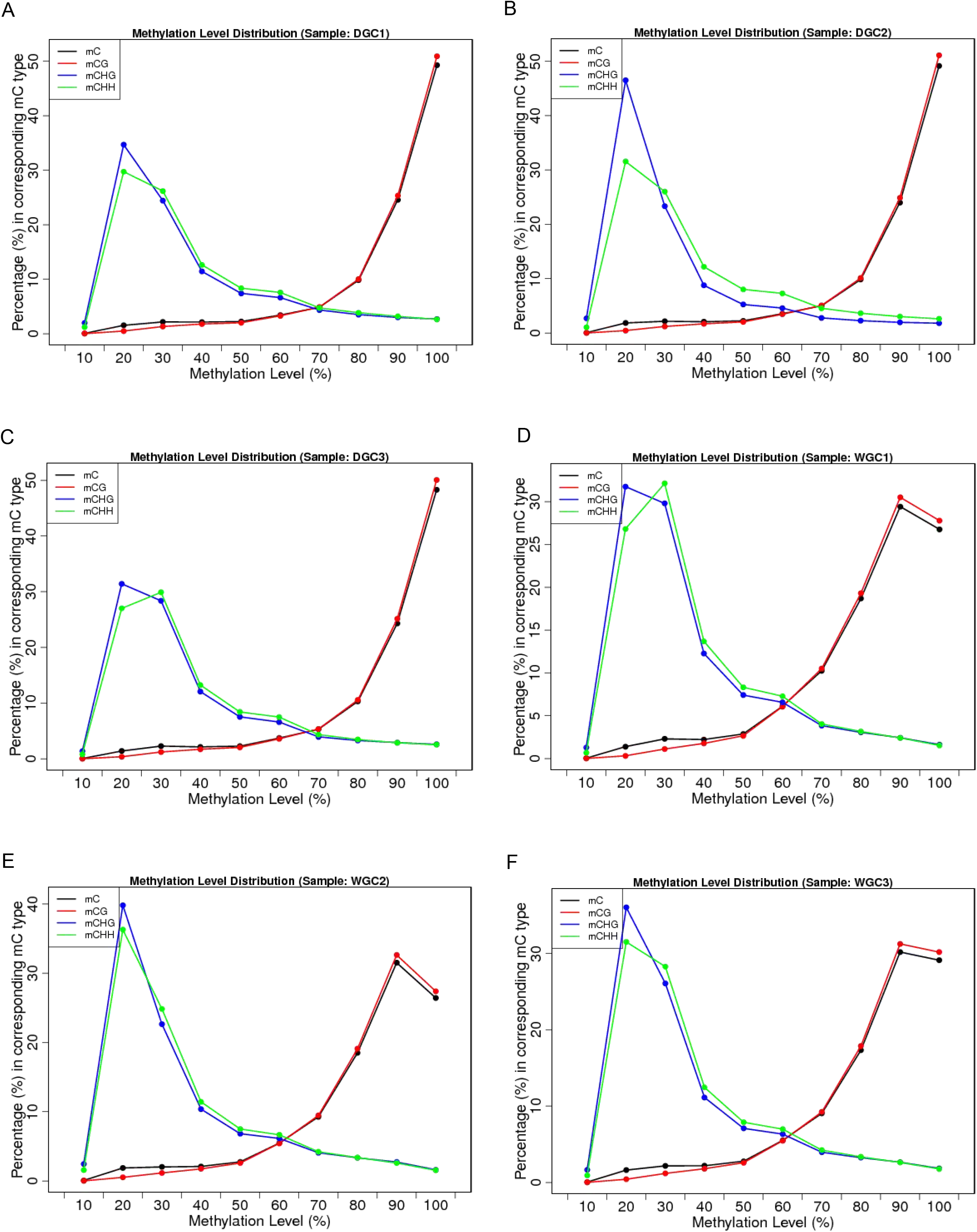
Distribution of methylation levels of mC in each sequence context. The x-axis was defined as the percentage of reads mC at a reference cytosine site. The y-axis indicated the fraction of total mC calculated within bins of 10%. DGC, domesticated grass carp; WGC, wild grass carp.

**Table 4.** Proportions of CG, CHG and CHH in all Methyl-cytosine

|  |  | mCG | mCHG | mCHH |
| --- | --- | --- | --- | --- |
| DGC1 | mC number | 26,890,172 | 208,311 | 750,652 |
|  | Proportions (%) | 96.557 | 0.748 | 2.695 |
| DGC2 | mC number | 26,953,558 | 323,600 | 784,048 |
|  | Proportions (%) | 96.053 | 1.153 | 2.794 |
| DGC3 | mC number | 26,520,365 | 219,138 | 811,594 |
|  | Proportions (%) | 96.259 | 0.795 | 2.946 |
| WGC1 | mC number | 26,700,829 | 228,299 | 848,145 |
|  | Proportions (%) | 96.125 | 0.822 | 3.053 |
| WGC2 | mC number | 27,258,571 | 220,189 | 831,299 |
|  | Proportions (%) | 96.286 | 0.778 | 2.936 |
| WGC3 | mC number | 26,934,911 | 222,172 | 821,714 |
|  | Proportions (%) | 96.269 | 0.794 | 2.937 |

Note: DGC, domesticated grass carp; WGC, wild grass carp. CG, CHG, and CHH (where H is A, C, or T).

### 3.4 DNA methylation levels of different genomic features

The heatmap presents the methylation landscape in different genomic features (whole genome, CGI, downstream 2 kb, upstream 2 kb, mRNA, repeat, CDS, exon), providing additional information as well as a global assessment of some of these components (Fig.2). CpG islands contained the highest numbers of CpG sites (approximately 10 - 20 CpG sites in a 200 bp window) compared with other genomic features. About 60% of CpG sites in CpG islands were hypermethylated (methylation levels >90%) in the heatmap (Figure2 and Supplementary Material Figure S2). The other genomic features generally contained 0 - 10 CpG sites in the 200 bp window. A lower proportion of CpG sites within whole genome, mRNA and repeat were hypomethylated (methylation levels < 10%) than CpG sites within remaining genomic features (CGI, downstream 2 kb, upstream 2 kb, CDS, exon).

**Fig.2.**
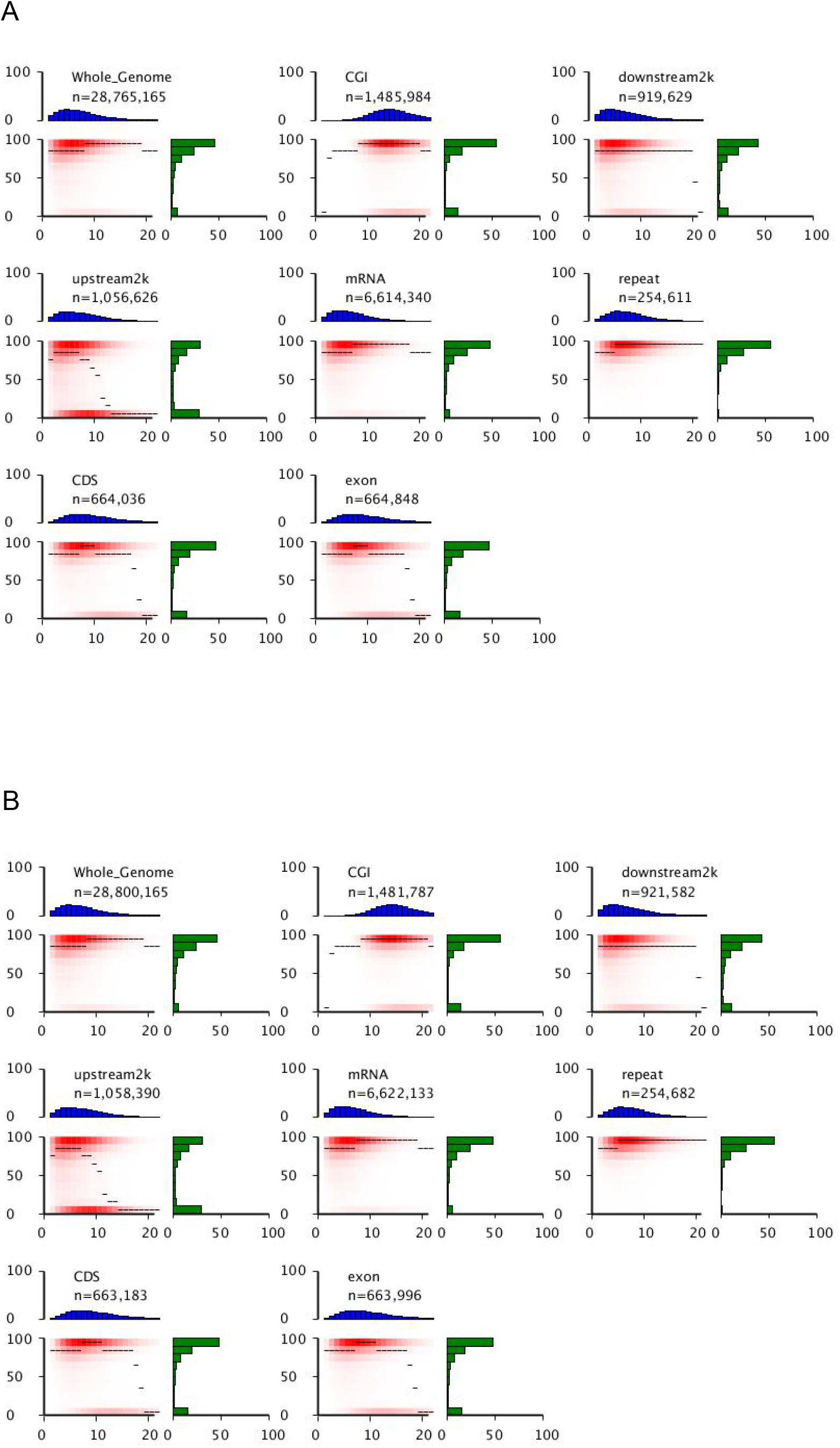
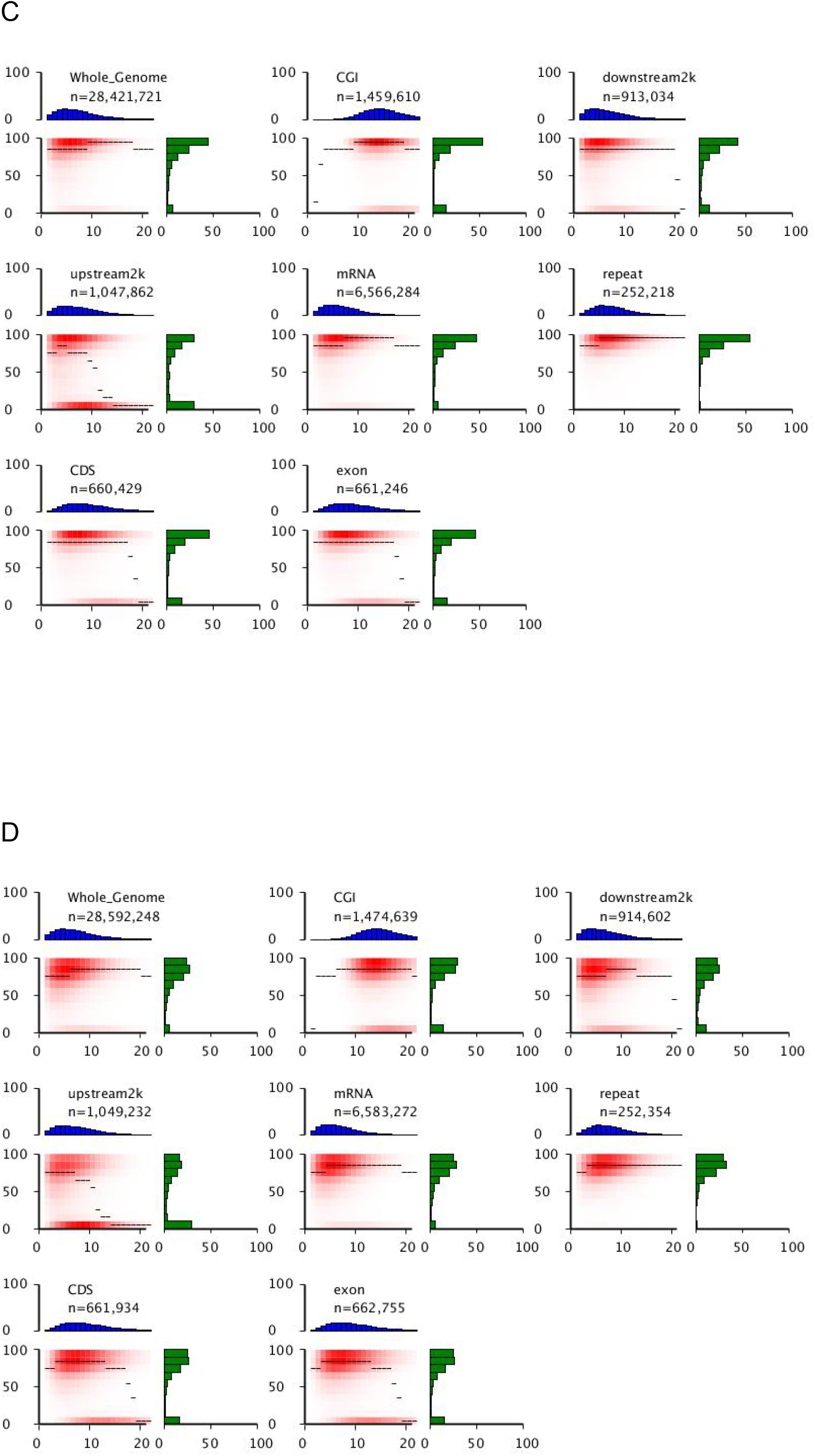
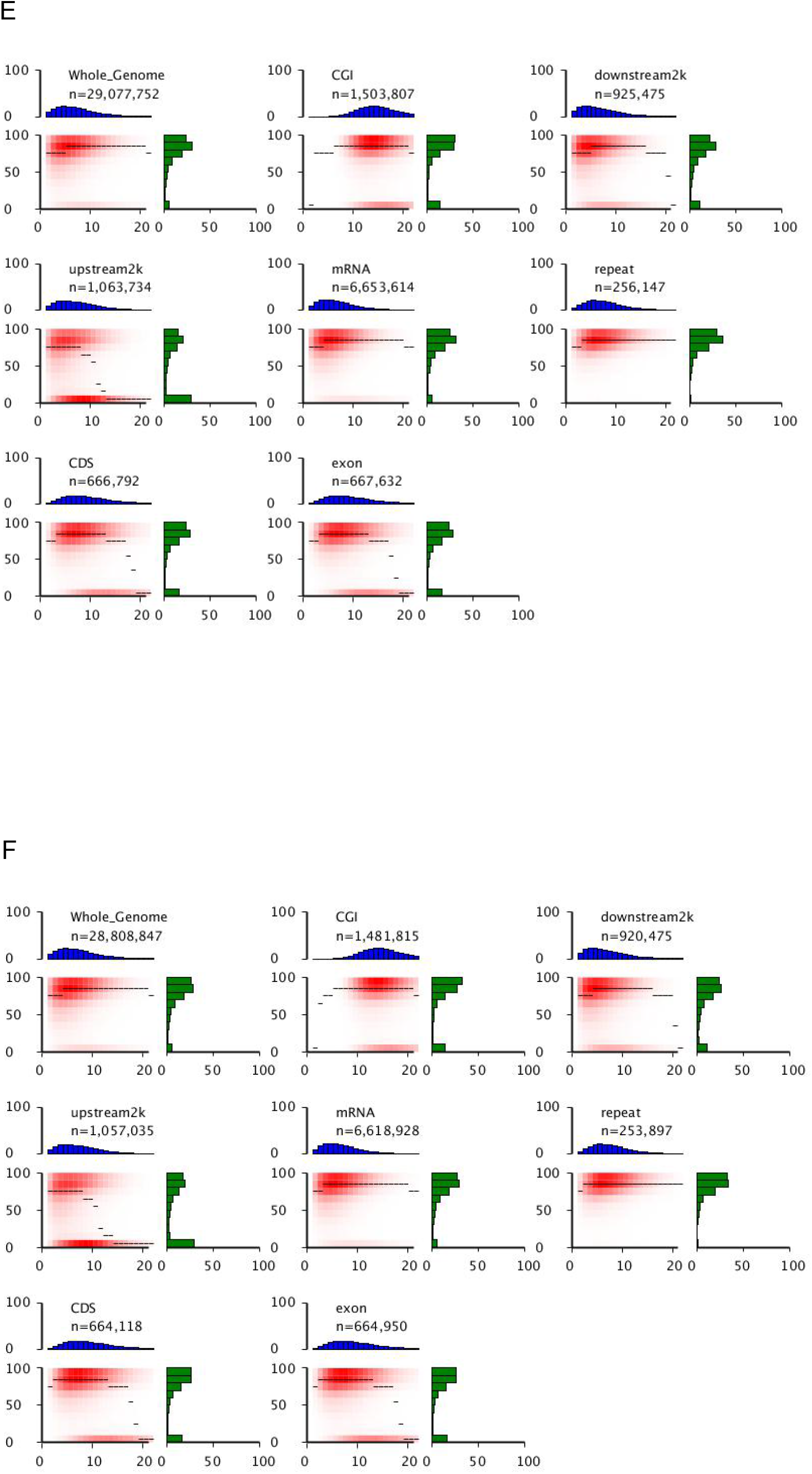
Heat maps of distinct methylation and CpG density patterns. CpG density (x-axis) was defined as numbers of CpG dinucleotides in 200bp windows. Methylation level (y-axis) was defined as average methylation level of cytosines in CpGs. The thin black lines within each heat map denoted the median methylation level of CpGs at the given local density. The red color gradient indicated abundance of CpGs that falled into bins of given methylation levels and CpG densities. The blue bar charts above each heat map showed the distribution of CpG densities, projected onto the x-axis of the heat maps. The green bar charts to the right of the heat maps show the distribution of methylation levels, projected onto the y-axis of the heat maps. (A) domesticated grass carp (DGC) sample 1; (B) sample DGC2; (C) sample DGC3; (D) wild grass carp (WGC) sample 1; (E) sample WGC2; (F) sample WGC3.

### 3.5 DNA methylation patterns across the entire transcriptional units at whole genome level

In order to reveal the relationship between DNA methylation profiles and genes expression in detail, Canonical DNA methylation profiles of the entire transcriptional units were divided into distinct functional elements to study the changes of methylation levels in different features (Fig.3). Methylation differences between CG and non-CpG methylation (CHG and CHH) are visible (Fig.3), as methylation levels of CG are higher than those of CHG and CHH across the entire transcriptional units. Another feature is a modest elevation in methylation level at internal exons and internal intron during transcriptional unit scanning. The lowest methylation level occurs in the first intron, followed by the first exon and downstream.

**Fig.3.**
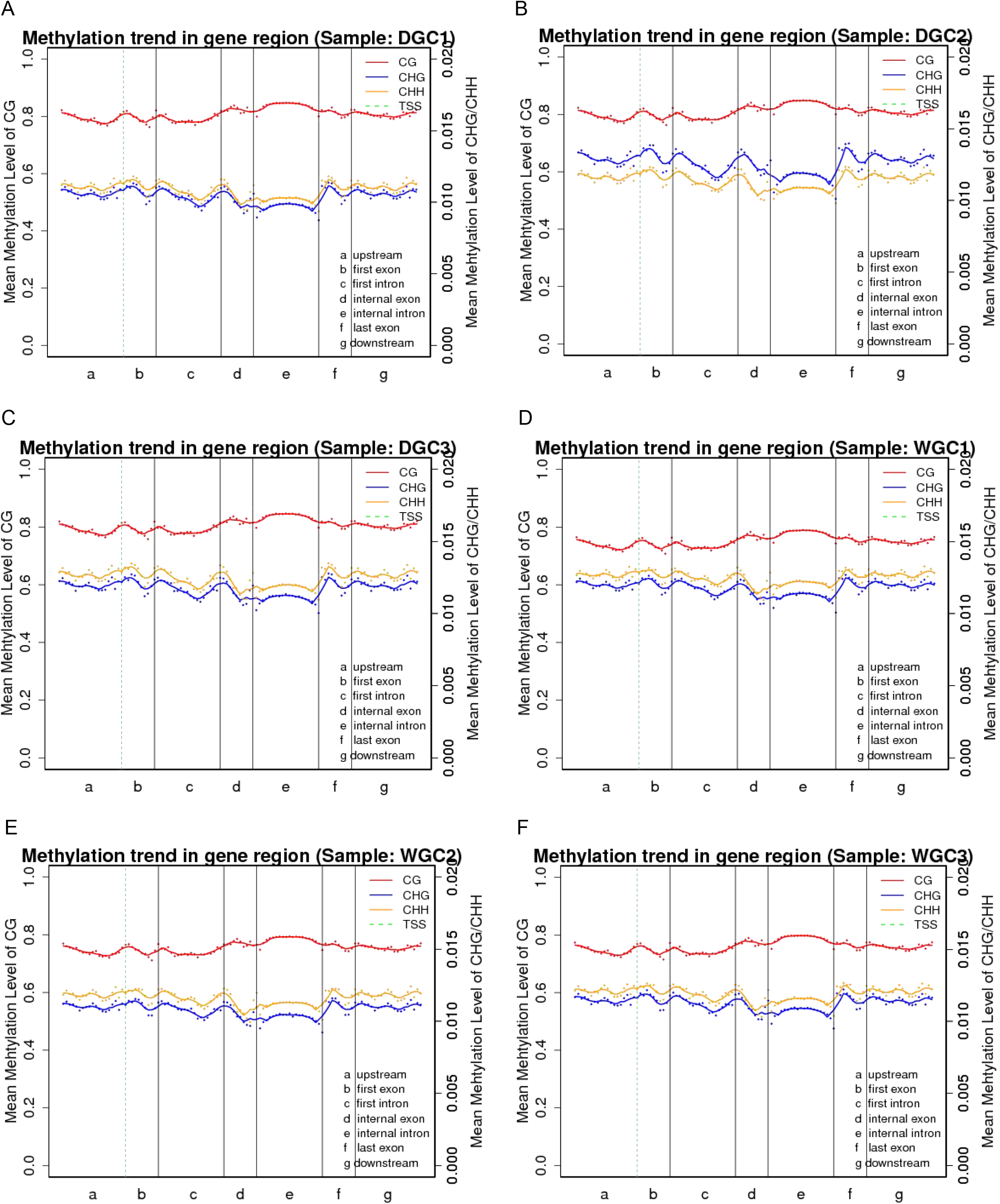
DNA methylation patterns across the entire transcriptional units at whole genome level. The canonical gene structure was defined by 7 different features, denoted by the x-axis. The length of each feature was normalized and divided into equal numbers of bins. Each dot denoted the mean methylation level per bin and the respective lines denoted the 5-bin moving average. Each feature was analyzed separately for the numbers listed in the table below the figure. The green vertical line indicated the mean location of the transcription start sites. DGC, domesticated grass carp; WGC, wild grass carp.

### 3.6 Identification and enrichment analysis of differential methylated regions

To characterize the differences of genome methylation levels between domesticated grass carp and wild grass carp groups, DMRs and differentially methylated genes (DMGs) were detected. For CG context methylation, a total of 533,235 DMRs were identified between the two groups, which corresponded to 20,010 DMGs in promoter regions and 27,016 DMGs in gene body.

GO enrichment analysis of DMGs was performed to provide significantly enriched GO terms corresponding to specific biological process, cellular component, and molecular function (Fig.4). In promoter regions, the over-represented GO terms in the biological process are cellular process, metabolic process, and biological regulation. The top enriched GO terms in the cellular component are cellular anatomical entity, intracellular and protein-containing complex. In terms of molecular function, the top enriched GO terms are binding, catalytic activity, and molecular transducer activity. In gene body region, the over-represented GO terms in the biological process are cellular process, metabolic process and biological regulation. The top enriched GO terms in the cellular component are cellular anatomical entity, intracellular and protein-containing complex. In terms of molecular function, the top enriched GO terms are binding, catalytic activity and transporter activity. KEGG pathway analysis, which is an alternative approach to categorize gene functions, was also conducted for the DMGs in promoter and gene body regions. In promoter region, DMGs were significantly enriched in pathways in cancer, neuroactive ligand-receptor interaction and PI3K-Akt signaling pathway. In gene body region, DMGs were significantly enriched in metabolic pathways, pathways in cancer and MAPK signaling pathway (Fig.5).

**Fig.4.**
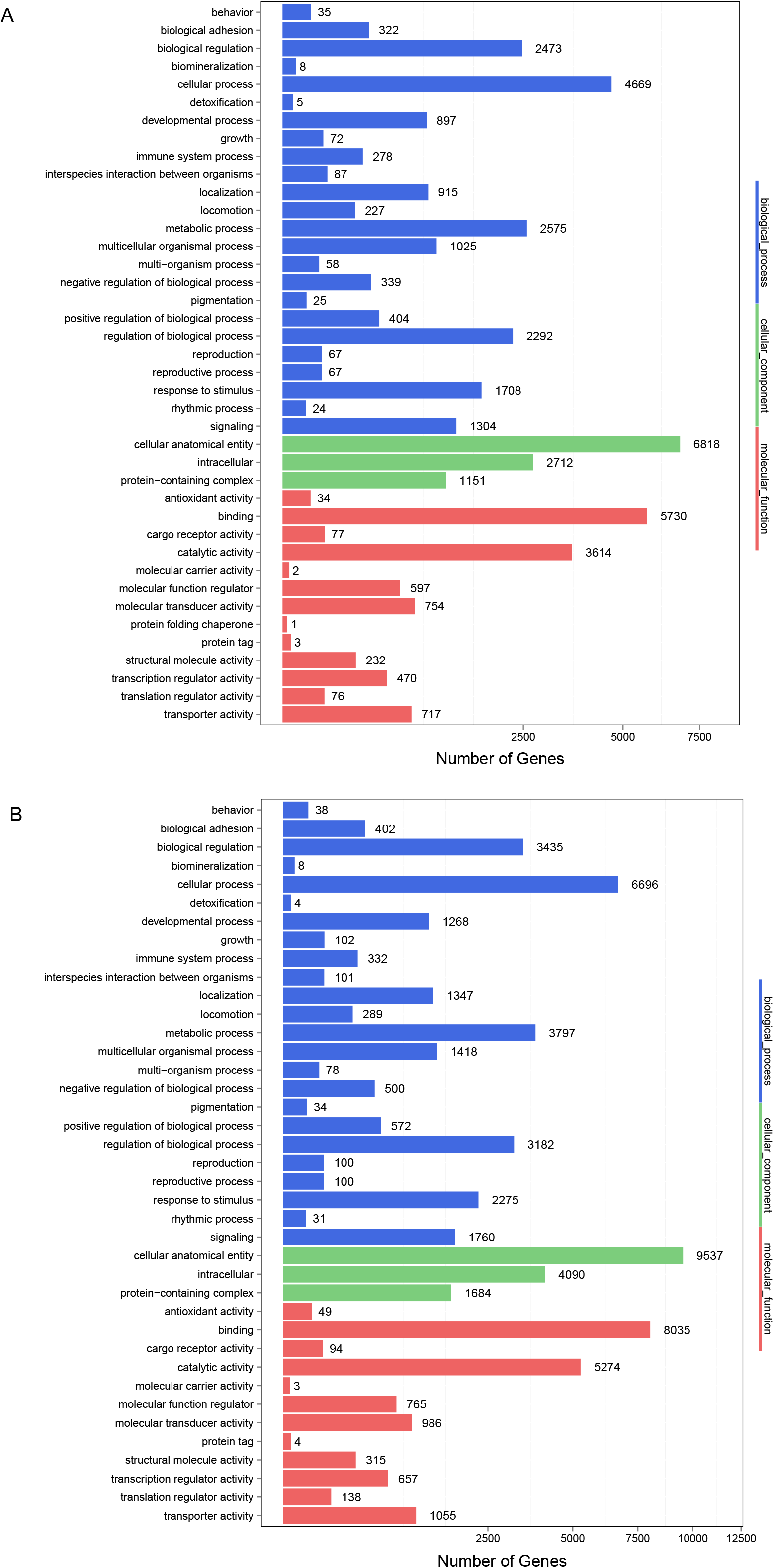
GO analysis of differentially methylated regions (DMRs)-related genes. The x-axis represented three domains of GO and the y-axis represented the gene number in each pathway and process. (A) GO analysis of DMRs-related genes in promoter region; (B) GO analysis of DMRs-related genes in gene body region.

**Fig.5.**
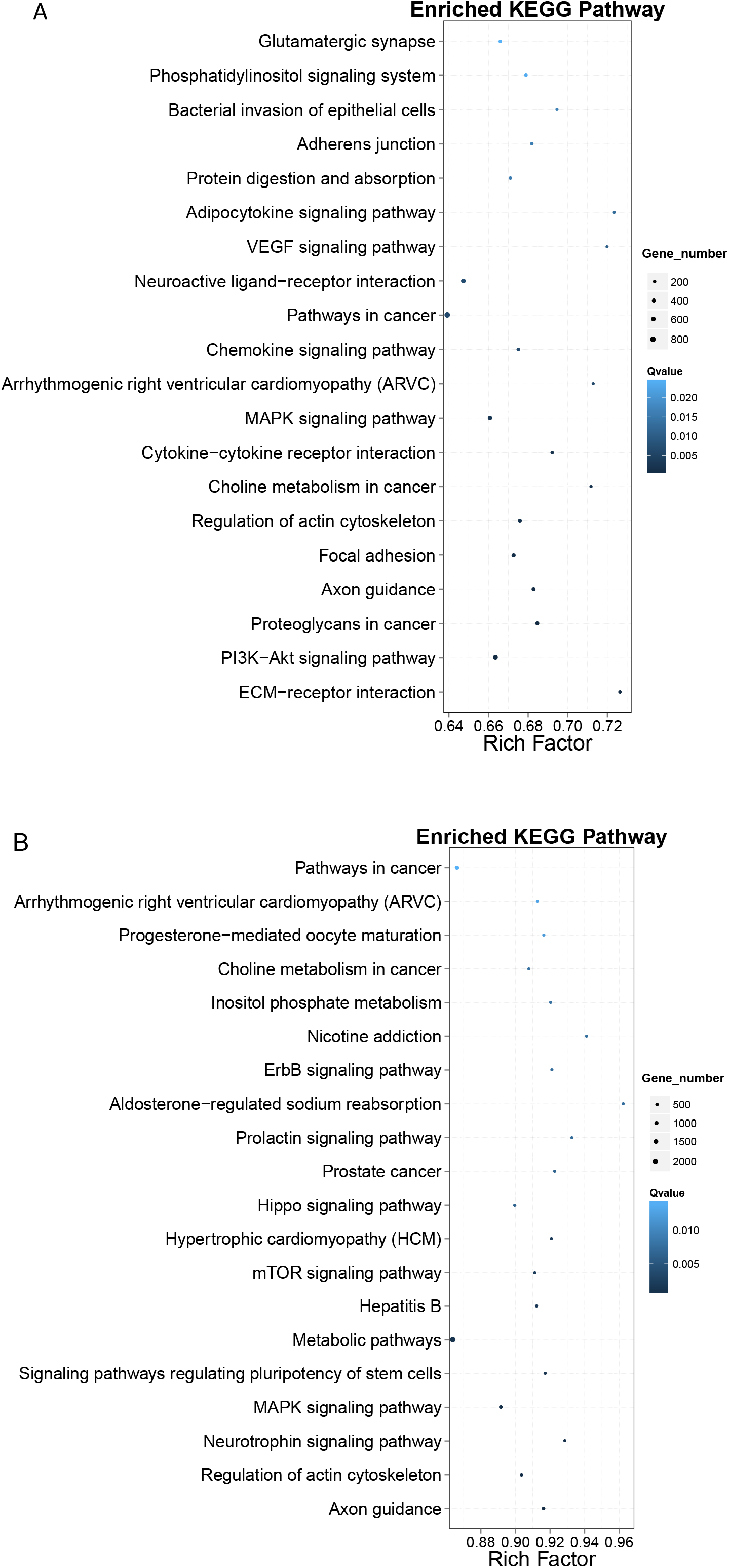
Pathway analysis of differentially methylated regions (DMRs)-related genes. The abscissa represented the richness factor, and the ordinate represented the enriched pathway terms. Q-value represented the corrected P, and a small Q-value indicated high significance. (A) Pathway analysis of DMRs-related genes in promoter region; (B) Pathway analysis of DMRs-related genes in gene body region.

### 3.7 Candidate DMGs associated with selection signatures

Many transcriptome and association studies have explored the molecular mechanisms underlying selection signatures in fish and other species, laying a foundation for our investigation of the involvement of DMGs in selection signatures. Besides, it is reported that DNA methylation, especially in the promoter regions, usually affects gene expression by different modes (Moore et al., 2012). Thus, 38 candidate DMGs in promoter region associated with the four selection signatures (growth and metabolism, immunity, foraging and learning behaviors), were identified according to the following criteria: (1) genes were differentially methylated in grass carp and wild grass carp groups; (2) genes were enriched in pathways related to selection signatures; (3) genes were differentially expressed in fish and other species or related with selection traits reported by previous studies. The 38 DMGs were used to construct the protein-protein interaction network using STRING database. We first obtained 107 pairs of interaction between 38 DEGs, and then these DMGs were functionally grouped according to KEGG pathway information (Fig.6). The network was divided into four parts, including foraging behaviors, learning behaviors, immunity and growth and metabolism. Among them, the network of learning behaviors was core and connected to the three other networks.

**Fig.6.**
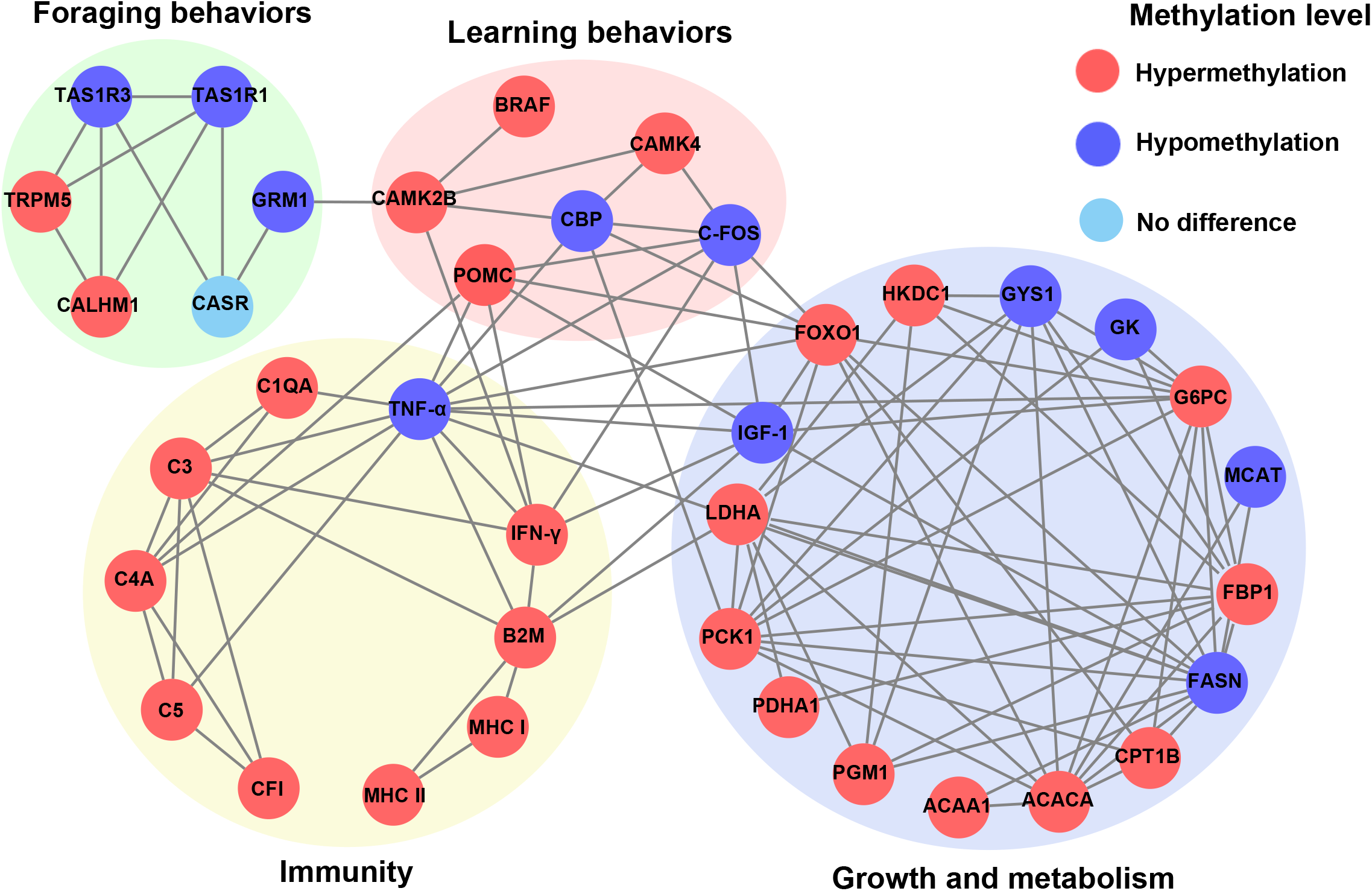
Protein-protein interaction networks of differentially methylated regions in different selection signatures.

## 4. Discussion

Domestication is a process by which humans select some phenotypes of wild animal species (i.e., morphological traits or growth). During the domestication process, some phenotypic traits of animal could be altered by the artificial selection to help domesticated animal species adapt to new environmental conditions (Sylvain et al., 2021). DNA methylation, one of the most important and stable epigenetic modifications in eukaryotes, can lead to heritable phenotypic and transcriptomic changes (Höglund et al., 2020). However, in aquatic species, there is still limited research that illustrate the effects of DNA methylation on selection signatures by comparing genome-wide methylation profiles between domesticated and wild fish species. Our study first systematically compared the genome-wide methylation profiles from domesticated and wild grass carp and identified some key differentially methylated genes related to selection signatures (growth and metabolism, immunity, foraging and learning behaviors) to uncover its genetic characters.

### 4.1 DNA methylation profiles in domesticated and wild grass carp

In the present study, we reported a genome-wide examination of DNA methylation in domesticated grass carp and wild grass carp. The mapping rates of clean reads ranged from 86.25% to 92.11% (Table 2), which is above the average value reported in other fish species (Pan et al., 2021). However, the WGBS results in this study were still consistent because bisulfite conversion reduces the complexity of the genomic sequence and the ability of most computational programs to align sequences onto the reference genome (Yang et al., 2020).

Further DNA methylation profile analysis was also conducted in our study across the distinct genomic features and entire transcriptional units. Our results showed that the genome-wide methylation patterns were similar between two groups. Approximately 9% of all total cytosines and 80% of CG were methylated and only a small proportions of methylation at non-CpG methylation (CHG and CHH) was observed within all regions, which is consistent with previous study (Cai et al., 2020). The high level of DNA methylation in CG content was a specific characteristic of animals. DNA methylation in CHH and CHG patterns is a major characteristic of plant methylomes and largely absent in animal methylomes (Zemach et al., 2010).

A modest elevation in methylation level at internal exons and intron was observed during transcriptional unit scanning. The lowest methylation level occurs in the first exons, followed by the first intron and last exon. These results was consistent with previous findings in grass carp (Cai et al., 2020), suggesting that mutagenic effects appear to occur at the first exons and internal exon and intron tend to be influenced by the regulatory impact of DNA methylation.

### 4.2 Key differentially methylated genes related to growth and metabolism

Domestication is a long process which forces animals to adapt to captivity by modifying an animal’ s growth and metabolism, immune response, foraging and social behaviors. In the process of domestication, fish undergo great changes in the growth and metabolism due to the difference of environmental conditions and food resource (Teletchea, 2016), which is closely associated with the differential expression of genes involved in growth and metabolism (Shen et al., 2021). In this study, several genes involved in growth and metabolism, such as IGF-1 (Insulin-like growth factors −1), GK (Glycerol kinase), GYS1 (Glycogen synthase 1), FASN (Fatty acid synthase, animal type), etc., exhibited hypomethylation in domesticated grass carp compared to wild grass carp. On the other hand, some genes related to growth and metabolism exhibited hypermethylation in domesticated grass carp, including G6PC (Glucose-6-phosphatase), PCK1 (Phosphoenolpyruvate carboxykinase 1), FBP1 (Fructose-1,6-bisphosphatase 1), FOXO1 (Forkhead box protein O1), ACAA1 (Acetyl-Coenzyme A acyltransferase 1) and CPT1 (Carnitine palmitoyltransferase I).

IGF-1, an important growth hormone, mediates the anabolic and linear growth and is thought to be a candidate gene for regulating muscle growth (Laron et al., 2001). GK is a key enzyme that catalyzes the first step in glycolysis (Enes et al., 2009). FOXO1, G6PC, FBP1 and PCK1 are key enzymes of gluconeogenesis (Lu et al., 2018). Our results indicated that, compared to wild grass carp, the glycolytic capacity was enhanced and the capacity for gluconeogenesis was inhibited in domesticated grass carp feeding commercial diet. These results was consistent with the previous findings that high-carbohydrate diets induce glycolysis and inhibit gluconeogenesis (Liu et al., 2021). GSY1 is a rate-limiting enzyme in glycogen synthesis and plays an important role in the synthesis of glycogen in the muscle (Martins et al., 2013). These results suggested that the capacity for muscle glycogen storage in domesticated grass carp possibly increased compared to that of wild grass carp.

FASN is believed to be the central enzyme of hepatic lipid accumulation (Dorn et al., 2010). ACAA1 and CPT1 play a certain role in the lipid degradation (Song et al., 2019). In the present study, our results suggested that, in domesticated grass carp, the ability to synthesize fat was enhanced and lipid degradation was inhibited compared to wild grass carp, which may account for elevated lipid levels of muscle in domesticated grass carp feeding commercial feed (Ashraf et al., 2011). According to the results, an explanation for the enhanced fat deposition in fish might be excessive energy intake in commercial diet (Wu et al., 2021). Overall, DNA methylation was likely to play an important role in regulating growth and metabolism of domesticated grass carp by influencing these gene expression.

### 4.3 Key differentially methylated genes related to immunity

To maximize profitability, domesticated fish were cultured in intensive farming conditions with limited space, high density and other stressors, which influence fish immune response negatively and even result in large-scale disease (Lin et al., 2018). It was reported that, in the process of domestication, the immune status of domesticated fish is affected negatively due to increased chronic stress by confinement, and makes fish more susceptible to pathogens and ultimately impair fish survival (Mandiki et al., 2011).

In this study, several immune related genes exhibited hypermethylation in domesticated grass carp compared to wild grass carp, including MHCI (MHC class I), MHCII (MHC class II), C1QA (Complement C1q subcomponent subunit A), C3 (Complement component 3), C4A (Complement component 4), C5 (Complement component 5), IFN-γ (Interferon gamma), etc. These genes suggested the expression of these genes might be downregulated. MHCI and MHCII are two cell surface proteins essential for the acquired immune system for antigen presentation to recognize foreign molecules in vertebrates and can promote the development and expansion of T cells (Huang et al., 2005). The complement system, an essential part of both innate and adaptive immunity in teleosts, is initiated by one or a combination of three pathways, the alternative, lectin and classical (Yang et al., 2016). IFN-γ, one of critical antiviral cytokines, modulate functions of the immune system by up-regulate major histocompatibility complex molecules (MHC I and MHC II) and directly activate other immune cells, such as macrophages and natural killer cells (Borden et al., 2007). In our results, these immune related genes mentioned above identified in domesticated grass carp exhibited hypermethylation, indicating a poor immune performance of grass carp during long-term domestication, which was consistent with the results of immune parameters that decreased levels of serum lysozyme, total protein in domesticated grass carp group reflected a reduction tendency in immune response (Lin et al., 2018).

### 4.4 Key differentially methylated genes related to foraging behaviors

During the domestication process, fish change their foraging habits to obtain food in captive conditions (Pasquet et al., 2018). In this study, foraging behaviors-related genes were also found to be under strong selection in domesticated carp. The genes of GRM1 (Metabotropic glutamate receptor 1), TAS1R1 (Taste receptor type 1 member 1) and TAS1R3 (Taste receptor type 1 member 3) exhibited hypomethylation in domesticated grass carp group compared to wild grass carp group, suggesting the expression of these genes might be upregulated. The TAS1R1 and TAS1R3 heterodimer receptor functions as an umami receptor, responding to L-amino acid binding, especially L-glutamate (Nelson et al., 2001). GRM1, which is widely expressed throughout the central nervous system and regulates synaptic signaling, is another L-glutamate receptor (Gabriel et al., 2009). Generally, these taste receptors play an important roles in perception of L-amino acids and feeding behavior (Cai et al., 2021). Thus, these genes with hypomethylation were likely to be upregulated in domesticated grass carp, facilitating the adaptation of domesticated grass carp to commercial feed by enhancing taste perception of nutrients in commercial diet in artificial rearing condition.

### 4.5 Key differentially methylated genes related to learning behaviors

Learning and memory enable the organism to plastically respond to the changing environment. Increasing research has investigated the learning (cognitive) and memory characteristics of fish in the past few decades, including spatial cognition, learned recognition, social learning, foraging activity, etc. (Dou et al., 2018). In this study, through domestication, social learning-related genes have also been under selection pressure. The gene POMC (Pro-opiomelanocortin) exhibited hypermethylation. In contrast, the genes of C-FOS (Proto-oncogene C-FOS) and CBP (CREB-binding protein) exhibited hypomethylation in domesticated grass carp group compared to wild grass carp group, suggesting the expression of these genes might be upregulated. C-FOS is necessary for consolidation of non-spatial hippocampal-dependent memory (Countryman et al., 2005), and C-FOS mRNA expressions are up-regulated in response to a variety of neuronal activation protocols, including behavioral training (Smulders et al., 2000) and long-term protentiation (Alberini et al., 2009). CBP is a coactivator of transcription that play an essential role in memory consolidation (Korzus et al., 2004). It was reported that gene expression of C-FOS and CBP facilitated the acquisition of novel feeding habits in fish through the memory formation (Peng et al., 2019; Dou et al., 2018). Besides, the C-FOS gene might be an important transcriptional factor to inhibit the expression of the anorexigenic gene POMC, resulting in an increase of the food intake of dead prey fish in mandarin fish (Peng et al., 2019). Therefore, it tempts us to speculate that the learning-related genes of C-FOS and CBP may be responsible for the acquisition of novel feeding habits by reinforcing memory in domesticated grass carp, which considerably increase the rate of domestication. Furthermore, individual food intake is enhanced through the interaction between the learning gene C-FOS and the appetite control gene POMC, contributing to the faster growth of domesticated grass carp indirectly.

## Declaration of competing interest

The authors declare that they have no known competing financial interests or personal relationships that could have appeared to influence the work reported in this paper.

## Acknowledgments

This study was funded by Central Public-interest Scientific Institution Basal Research Fund (CAFS, NO. 2021XT03) and China Agriculture Research System of MOF and MARA (No.CARS-45-21).

## Notes

### Competing Interest Statement

The authors have declared no competing interest.

### Summary of Updates

Tables revised; Introduction revised; Materials and methods revised; Results revised.

